# Inhibition of the Platelet-activating factor receptor for the treatment of Amyotrophic Lateral Sclerosis?

**DOI:** 10.1101/224030

**Authors:** Marcelo R. S. Briones, Amanda M. Snyder, Elizabeth B. Neely, James R. Connor, James R. Broach

**Affiliations:** Department of Health Informatics, Escola Paulista de Medicina, UNIFESP, Brazil, Rua Pedro de Toledo 669 L4E, São Paulo, SP, 04039-032; Department of Biochemistry, Institute for Personalized Medicine, Penn State College of Medicine, 500 University Drive, Hershey PA 17033; Department of Neurosurgery, Penn State College of Medicine, 500 University Drive, Hershey PA 17033

**Keywords:** Amyotrophic Lateral Sclerosis, Platelet-activating factor, Antagonists

## Abstract

Cerebrospinal Fluids (CSF) of Amyotrophic Lateral Sclerosis (ALS) patients have increased levels of the inflammatory cytokine IL-18. Because IL-18 is produced by dendritic cells stimulated by the Platelet-activating factor (PAF), a major neuroinflammatory mediator, it is expected that PAF is involved in ALS. Pilot experimental data on amplification of PAF receptor (PAFR) mRNA by RT-PCR show that PAFR is overexpressed, as compared to age matched controls, in the spinal cords of transgenic ALS mouse model SOD1-G93A, suggesting PAF mediation. Although anti-inflammatory drugs have been tested for ALS before, no clinical trial has been conducted using PAFR specific inhibitors. Therefore, we hypothesize that administration of PAFR inhibitors, such as Ginkgolide B, PCA 4248 and WEB 2086, have potential to function as a novel therapy for ALS, particularly in SOD1 familial ALS forms. Because currently there are only two approved drugs with modest effectiveness for ALS therapy, a search for novel drugs and targets is essential.

## Introduction

Amyotrophic Lateral Sclerosis (ALS), a Motor Neuron Disease, is the third most prevalent neurodegenerative disease (4 cases per 100,000 people), being Alzheimer’s disease and Parkinson’s disease the first and second, respectively (1). ALS is a progressive, irreversible and fatal neurodegenerative disease for which no effective therapy exists. Riluzole was for 22 years the only drug approved in ALS but only confers a modest improvement in survival although providing relief of respiratory symptoms (2,3). Edaravone was approved in 2017 but its therapeutic effectiveness has not been fully established (4). ALS affects upper and lower motor neurons with pronounced degeneration of Alpha motor neurons that innervate extrafusal fibers of skeletal muscle (5). The clinical manifestations of ALS are muscle atrophy, dysphagia, dysarthria, spasticity, hyperreflexia, Babinski’s sign, fasciculation and respiratory failure. Previous observations have established that only 10% of ALS patients have a family history of the disease, which means that 90% of patients have no near relatives who have presented with the disease. Twin studies have estimated ALS heritability to be 60-70% (6), suggesting that many patients who present with sporadic ALS (SALS) may also have an underlying genetic cause. C9orf72 and SOD1 are considered the ‘major’ ALS-causing genes. Mutations in C9orf72 are observed in 38% FALS and 8% SALS while mutations in SOD1 have been reported in 13% FALS and about 1% SALS (7,8). Since SOD1 has been the first ALS-causing gene described, in 1993, transgenic mouse models have been well established for the SOD1 ALS and have been used in several studies bearing on the basic mechanisms of ALS. SOD1 ALS is caused by genetic gain of function and the mouse model has the G93A substitution, the most widely used transgenic mouse lineage (9).

Several alterations in brain chemistry are associated with ALS ranging from glutamate imbalance in upper motor neuron synapses, inflammation and astrocyte activation. Despite its demonstrated role in other neurological disorders (10), PAF, a major neuroinflammatory mediator, has not been characterized in ALS. The synthesis of PAF occurs either via remodeling (lyso-PC acetyltransferases, LPCAT 1 and 2), or by *de novo* synthesis via phosphocholinetransferase (PAF-PCT) (11). Analysis of Cerebrospinal Fluids (CSF) of ALS patients show increased levels of inflammatory cytokines, particularly IL-18, also known as interferon-gamma inducing factor (12). IL-18 is produced by dendritic cells stimulated by PAF (13) which suggests that PAF might be associated with ALS. Clinical tests so far conducted have not included PAFR specific inhibitors (14). PAFR inhibitors are used as therapy for allergies, cancer and cardiac disease (15) and could be repurposed for ALS treatment.

## Hypothesis

We hypothesize that administration of Platelet-activating factor receptor inhibitors has potential to function as a novel therapy for ALS, particularly in SOD1-familial forms.

## Evaluation of the hypothesis

The involvement of inflammation in the pathogenesis of ALS is increasingly recognized but not well understood yet. It has been shown that among all inflammation-related IL-1 family cytokines (IL-1β, IL-18, IL-33, IL-37) and their endogenous inhibitors (IL-1Ra, sIL-1R2, IL-18BP, sIL-1R4) only IL-18 and its endogenous inhibitor, IL-18BP, are significantly increased in CSF of patients with ALS as measured by the ELISA method (12). The increase of total free IL-18 suggests the activation of IL-18-cleaving inflammasome. Whether IL-18 upregulation in ALS patients is a consequence of inflammation or one of the causes of the pathology still needs to be tested. PAF is a very important mediator of inflammatory response. Also known as PAF-acether or AGEPC (acetyl-glyceryl-ether-phosphorylcholine) is a potent phospholipid activator and mediator of several leukocyte functions, platelet aggregation and degranulation, inflammation, and anaphylaxis. It is also involved in changes to chemotaxis of leukocytes, vascular permeability, oxidative burst, and increased arachidonic acid metabolism in phagocytes. High PAF levels are associated with a variety of medical conditions including: allergic reactions, multiple sclerosis, stroke, myocardial infarction, colitis, inflammation of the large intestine and sepsis.

PAF is produced by several cell types, especially those involved in host immunity, such as platelets, macrophages, neutrophils, monocytes and endothelial cells. PAF is constitutively produced in low levels by these cells and its synthesis is controlled by PAF acetylhydrolases activity. In response to specific stimuli it is produced in larger quantities by inflammatory cells (16). PAF acts on a specific receptor, PAFR, expressed in mature and immature dendritic cells (17). Because PAF induces monocyte-derived dendritic cells but not macrophages to secrete IL-12 and IL-18 (13), it is expected that PAF, and it receptor, are involved in ALS as the source of increase IL-18 in ALS patients might be PAF activated dendritic cells. This is the background of the hypothesis. Therefore the upregulation of PAFR in ALS patients, or the ALS-SOD1 mouse experimental model, might be suggestive of a PAF role in ALS.

## Empirical data

A pilot study data on upregulation of PAFR in the ALS experimental mouse model is presented. All animal experiments were performed in accordance with protocol #46763 approved by the IACUC of Pennsylvania State University. Experimental mice (SOD1-G93A strain, Jackson Laboratories) and control mice (C57BL/6J, Jackson Laboratories) were sacrificed at 110 days of age (symptomatic). Experimental mice in the symptomatic group are not severely affected at that time point and do retain normal feeding and grooming behavior although locomotion is affected, as evaluated by the rotarod performance test. Animals were deeply anesthetized using KAX (100mg/kg ketamine, 10mg/kg xylazine and 3mg/kg acepromazine, to be injected i.p. at a weight-adjusted dose of 0.1 mL / 10g bodyweight). The level of anesthesia was assessed by lack of response to toe and tail pinch, followed by cardiac puncture to remove blood and then exsanguination. Mice were decapitated immediately following exsanguination with sharpened scissors. Spinal cord tissue was removed and frozen in liquid nitrogen.

Lumbar sections of the spinal cord were dissected and RNA isolated using the DNA/RNA extraction kit (DNAEssy, Qiagen) and quantitated using Nanodrop (ThermoFisher Scientific). Three ALS mice and three age matched controls were used in RT-PCR with SuperScript IV reverse transcriptase and PCR kit. PCR was carried with PAFR specific primers (87F-5’-GGT GACTT GGCAGT GCTTT G and 530R-5’-CACGTTGCACAGGAAGTTGG) located in two different exons (positions 87 in exon 1 and position 15,510 in exon 2 of PAFR gene). For RNA load control amplification of 18S rRNA was used (primers 757F-5’-CCCCTCGAT GCT CTT AGCT G and 1,516R-51-CCCGGACATCTAAGGGCATC). In Fig. 1 the amplification of PAFR in ALS mice and controls is shown. All three ALS-SOD1 mice (A1, A2 and A3) show higher PAFR expression as compared to controls (C1, C2 and C3).

**Fig 1.**
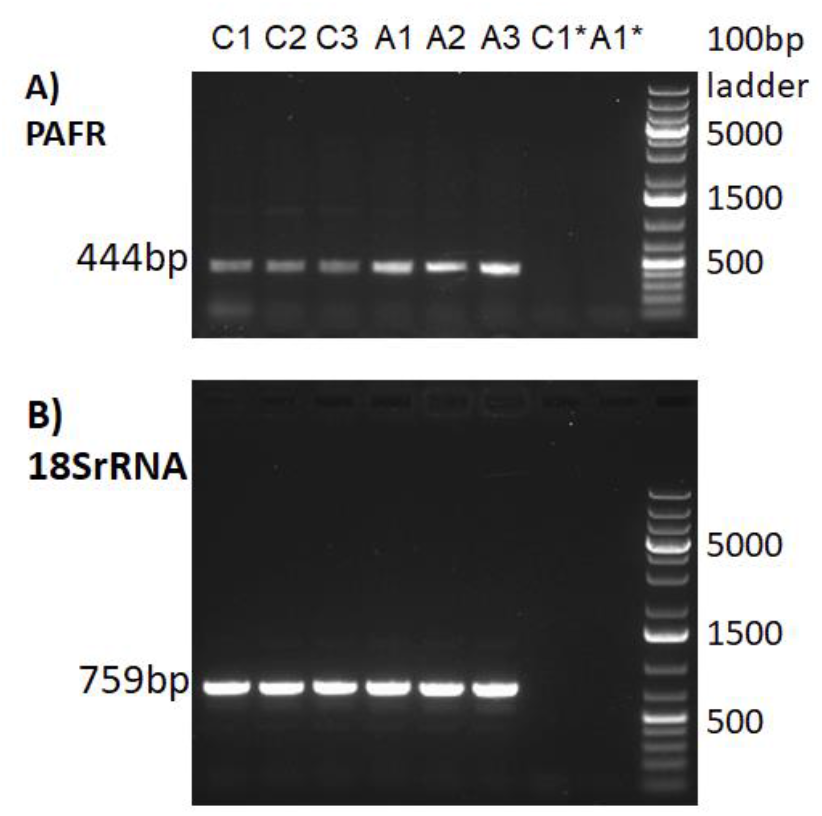
RT-PCR of PAFR mRNA. In (A) amplification with PAFR specific primers in ALS/SOD1-G93A mice (A1, A2 and A3) and in BL6 controls (C1, C2 and C3). In (B) amplification of 18S rRNA for RNA load control in the same samples as in (A). PAFR yields a 444bp amplicon and 18SrRNA a 759bp amplicon. Primer annealing temperatures are 62°C for PAFR and 63°C for 18SrNA. C1* and A1* indicate RT-PCR controls without reverse transcriptase to show that amplicons are entirely dependent on reverse transcriptase activity and therefore not due to DNA contamination in RNA preparations.

Gel densitometry analysis of the gel electrophoresis indicates that PAFR is in average 1.6 times overexpressed in ALS-SOD1 mice as compared to BL6 mice controls (Table 1).

**Table 1.** Densitometry of gel electrophoresis depicted in Fig. 1. Quantitation of specific gel band area by ImageJ software (https://imaqei.nih.qov/ij/. A1, A2 and A3 are ALS/SOD1 mice and in C1, C2 and C3 the BL6 non ALS controls.

| Mouse | Area (pixels) | Mean | Min | Max |
| --- | --- | --- | --- | --- |
| C1 | 330 | 75.003 | 31 | 140 |
| C2 | 330 | 72.209 | 34 | 136 |
| C3 | 330 | 71.718 | 36 | 122 |
| A1 | 330 | 122.258 | 37 | 221 |
| A2 | 330 | 108.527 | 39 | 251 |
| A3 | 330 | 127.279 | 37 | 252 |

## Future experiments required

The upregulation of PAFR in CNS of ALS mice was tested in the lumbar section of the spinal cord. However ALS affects glutamate synapses in the motor cortex. Lower motor neurons have acetylcholine synapses. Therefore quantitative RT-PCR (18) should be performed not only in spinal cord RNA but also in motor cortex in the brain. More refined quantitation using quantitative PCR (19) should be performed to guarantee that the comparative expression is in the linear range of amplification reaction and not underestimated by signal saturation. Also, upregulation of PAFR peptide in the motor cortex and spinal cords should be tested with anti-PAFR antibodies to corroborate results obtained with mRNA upregulation.

The effect of PAFR inhibitors can be tested initially in ALS-SOD1 mice by oral administration of Ginkolide B with dosage appropriately scaled for mice. The effects on ALS symptoms, such as hind limb impairment can be tested against age matched controls.

## Consequences of the hypothesis and discussion

Only two FDA approved drugs, Riluzole and Edaravone, are available currently for ALS therapy. Riluzole modulates glutamate neurotransmission by inhibiting both glutamate release and postsynaptic glutamate receptor signaling. Edaravone is a free radical scavenger (20). Riluzole increases patient survival by 3 to 6 months and relieves respiratory discomfort while the therapeutic effects of Edaravone are still controversial (21). A therapeutic possibility involving PAF antagonists via PAFR inhibition is discussed here. The rationale is that the augmented IL-18 in CSF in ALS patients is consistent with pilot experimental data indicating the overexpression of PAFR in SOD1 ALS mouse model. Several natural and synthetic PAF inhibitors are known, used with therapeutic purposes and can be tested for relief or cessation of ALS symptoms in the animal model. If effective these compounds might prove a valuable tool in ALS therapy. The ALS mouse model used in pilot experiment is the SOD1-G93A gain of function model, therefore it can be speculated that PAF inhibitors might have an effect at least for the treatment of the SOD1 ALS subtype which corresponds to 13% of FALS and about 1% of SALS.

## Ethics Statement

All animal experiments were performed in accordance with protocol #46763 approved by the IACUC of Pennsylvania State University.

## Author Contributions

MRSB conceived the hypothesis, planned performed the experiments and wrote the manuscript. AMS planned, performed the experiments and edited the manuscript. EBN performed the experiments. JRC and JRB discussed the results and edited the manuscript.

